# Shear Stress Regulates ABCA1-dependent Membrane Cholesterol Content in Endothelial Cells Facilitating H2S-dependent Vasodilation

**DOI:** 10.1101/2025.04.21.649801

**Authors:** Jacob R. Anderson, Nancy L. Kanagy, Laura V. Gonzalez Bosc, Jay S. Naik

## Abstract

Endothelial cells (ECs) express an array of integral membrane proteins, including ion channels and transporters that contribute to blood flow regulation and cell-cell communication. Many of these membrane proteins are regulated by plasma membrane cholesterol content. The ATP-binding cassette family a1 (ABCA1) transporter is a regulator of membrane cholesterol content. We have shown increased ABCA1 mRNA expression and reduced EC membrane cholesterol in resistance mesenteric arteries compared to conduit arteries. Previous studies suggest shear stress (SS) can increase or decrease ABCA1 expression in a cell-type-dependent manner.

**Hypothesis:** SS sustains lower EC membrane cholesterol concentration through ABCA1-mediated cholesterol transport, facilitating H_2_S-mediated vasodilation.

**Methods:** The effect of SS on ABCA1 and membrane cholesterol content was assessed in pressurized mesenteric arteries from male Sprague-Dawley rats and cultured human aortic endothelial cells. Pressure myography was used to assess the effects of ABCA1 inhibition on H_2_S-mediated vasodilation. Filipin was used to assess EC membrane cholesterol content.

**Results:** SS increased ABCA1 expression in the endothelium of mesenteric arteries and cultured human aortic endothelial cells and markedly reduced EC membrane cholesterol. Inhibition of ABCA1 increased EC membrane cholesterol content and abolished H_2_S-induced vasodilation.

**Conclusion:** SS facilitation of EC-dependent vasodilation appears to be mediated by membrane cholesterol content.

## Introduction

The endothelium comprises a monolayer of highly specialized endothelial cells (ECs) lining the luminal surface of all blood vessels and occupies a fundamental position in the cardiovascular system. The endothelium regulates complex intercellular interactions, including vascular tone, vascular permeability, and innate immune response^1,2^. The endothelium senses and integrates various stimuli to regulate vascular tone through the synthesis and release of vasoactive gas molecules, such as nitric oxide, carbon monoxide, and hydrogen sulfide (H_2_S) ^3^.

H_2_S has been reported to be an endothelial-derived hyperpolarization factor contributing to vasodilation^4^ and exerting antioxidant and anti-inflammatory protective effects on the cardiovascular system^5^. The enzymatic synthesis of H_2_S involves cystathionine-γ-lyase, cystathionine-β-synthase, and 3-mercaptopyruvate sulfurtransferase; the specific enzymatic pathways vary across vascular beds, adding complexity to the regulation of H_2_S-mediated vasodilation^6,7^. Our previous work has shown that H_2_S sensitivity in conduit arteries (>300 μm) is dependent on the content of membrane cholesterol in ECs, whereas resistance arteries (<150 μm) are inherently sensitive to H_2_S^8^.

Membrane cholesterol is a fundamental constituent of the lipid bilayer, impacting membrane fluidity, stability, and organization^9^. Interestingly, our lab has shown that native EC membrane cholesterol content is greater in EC from conduit arteries than in those from resistance arteries,^8^ suggesting innate regional differences in membrane cholesterol content along the vascular tree. In addition, our single-cell RNAseq analysis supported by immunohistochemistry indicates that ECs from resistance arteries express greater levels of the reverse cholesterol transport genes ATP-binding cassette protein A1 (ABCA1) and phospholipid transfer protein (PLTP) compared to ECs from conduit vessels^10^. The ATP-binding cassette (ABC) transport proteins, a family of eight genes, are responsible for transporting various hydrophobic compounds across membranes. ABCA1 specifically facilitates membrane cholesterol efflux through a process called reverse cholesterol transport in multiple cell types^11,12^. Taken together, these findings suggest that reduced membrane cholesterol within resistance arteries enhances endothelial-mediated vasodilation, supporting the postulate that membrane cholesterol is an important mediator of EC function. However, despite the observed regional heterogeneity in ABCA1 expression within the vascular tree, it is unknown whether ABCA1 mediates innate differences in EC cholesterol content. Additionally, the mechanism mediating the differential ABCA1 expression between conduit and resistance arteries also remains unknown. Previous studies suggest shear stress (SS) can both increase and decrease ABCA1 expression, potentially in a cell-type-dependent manner^13–15^.

Vascular SS is a mechanical force resulting from friction against the endothelium generated by blood flowing along its surface. Vascular ECs sense SS through the activation of mechanosensors, translating it into biochemical signals influencing cell morphology, vascular tone, and gene expression^16,17^. As the luminal diameter of an artery decreases, SS exponentially increases^18^, suggesting that differences in SS may mediate cellular heterogeneity along the vascular tree. Therefore, we hypothesized that SS sustains lower EC membrane cholesterol concentration through ABCA1-mediated cholesterol transport, facilitating H_2_S-mediated vasodilation.

## Methods

### Cell Culture

Primary HAoECs (Cell Applications S304-05a, P4-P6) were used for all cell culture experiments. Cells were plated on cell culture dishes coated with 0.1% Fibronectin (Sigma, FC010) and maintained at 37°C, 5% CO_2_ in endothelial growth media V2 (Cell Applications). To expose cells to SS, ECs were seeded on μ-Slide 0.4 Luer ibiTreat (ibidi 80176). Unidirectional, recirculating, laminar flow was applied to confluent monolayers using a peristaltic pump [6 dyne/cm^2^, 20 dyne/cm^2^, or 0 dyne/cm^2^ for static control] with an inline wind Kessel (to dampen pulsatile oscillations) and bubble trap.

### Animals

Sprague-Dawley rats (Envigo, 250–300 g, 9–10 wk. old) were used for all experiments. Rats were euthanized with a lethal concentration of pentobarbital sodium (200 mg/kg I.P.), and mesenteric arteries were collected for experiments. The Institutional Animal Care and Use Committee of the University of New Mexico School of Medicine reviewed and approved all animal protocols. All protocols conformed to National Institutes of Health guidelines for animal use.

### Preparation of Endothelial Tubes

The mesenteric arcade was removed and immediately placed in ice-cold HEPES buffered physiological saline solution (HPSS; in mM: 130 NaCl, 4 KCl, 1.2 MgSO_4_, 4 NaHCO_3_, 1.8 CaCl_2_, 10 HEPES, 1.18 KH_2_PO_4_, and 6 glucose; pH adjusted to 7.4 with NaOH), containing 10 μM sodium nitroprusside and 0.1% bovine serum albumin (BSA). Veins were removed, and arteries were cleared of connective and adipose tissue. Arteries (50-150μm) were flushed of blood and pretreated with probucol (100 μM in 2% pluronic acid) or vehicle at 37°C for 1 hour in HPSS containing 10% FBS and 0.1% bovine serum albumin. Endothelial tubes were isolated as previously described^19^. Briefly, arteries were then placed in an HPSS digestion solution (10% FBS, 0.1% bovine serum albumin, 0.62 mg/mL of papain, 1.5 mg/ mL of collagenase II, and 1 mg/mL dithiothreitol) with continued treatment with probucol or vehicle. The arteries were incubated in the digestion solution for 33 minutes at 37°C. The digestion solution was replaced with ice-cold HEPES buffered physiological saline solution (HPSS; in mM: 130 NaCl, 4 KCl, 1.2 MgSO_4_, 4 NaHCO_3_, 1.8 CaCl_2_, 10 HEPES, 1.18 KH_2_PO_4_, and 6 glucose; pH adjusted to 7.4 with NaOH), 10% FBS and 0.1% BSA and artery segments were gently triturated to disrupt the adventitia and smooth muscle layers. Isolated EC tubes are then transferred using a transfer pipette and allowed to adhere to a glass slide. A pap-pen was used to create a hydrophobic pocket on the slide with sufficient surface tension to hold the EC tubes, antibody, and other solutions within a confined area. A Hamilton 250μL glass syringe was used to apply staining solutions and perform washes. EC tubes were mounted with ProLong Gold antifade mounting reagent.

### Quantification of Membrane Cholesterol Content

Isolated endothelial tubes were plated on glass coverslips and fixed with 4% paraformaldehyde (20 min at RT). Endothelial cell membrane cholesterol was detected by incubating cells with the fluorescent cholesterol marker filipin III (100 μg/mL) for 45 minutes at 37 °C under light-protected conditions. Coverslips were then mounted on slides using Prolong Gold Antifade (Thermo Fisher P10144). Samples were imaged using a fluorescent confocal microscope (Zeiss LSM 800 AxioObserver.Z1/7; Gottingen, Germany) with a 353 nm laser (excitation) and a 465 nm long-pass filter (emission), combined with a Plan-Apochromat 63/1.4 numerical aperture (NA) oil immersion objective. A negative control lacking filipin was used to control for background signal. Filipin staining was quantified using ImageJ^20^. A total of 4 tubes/artery from five animals were analyzed. The mean fluorescence intensity of each endothelial tube was measured and averaged to determine a mean fluorescence value for each animal.

HAoECs were stained as previously described above with filipin, and the samples were imaged using a Zeiss LSM800. Images contained 8-10 images with 7-10 cells per image for three independent experiments. Regions of interest were created at approximately 1 mm from the membrane, both intra- and extracellularly, to isolate the membrane from the intracellular compartments. Images were collected within one optical section showing membrane max diameter to ensure the focus was on the cell center, and the mean fluorescence intensity per cell was measured and averaged to determine cholesterol content.

### Quantification of ABCA1 Protein Expression

Endothelial ABCA1 expression was quantified using immunofluorescence in *en-face* small arteries and HAoECs. Small (<150 μm) were cannulated and pressurized^21^. Briefly, pressurized mesenteric arteries were fixed with 4% paraformaldehyde for 20 minutes at room temperature. Samples were blocked with 5% non-donkey serum (NDS) and treated with ABCA1 primary antibody (1:100) (Thermo Fisher, PA1-16789) overnight at 4°C. Endothelial tubes were then blocked with 5% NDS for 1 h at 37°C and incubated with donkey-α-rabbit 488 (1:1000) (Jackson ImmunoResearch Labs, 711-546-152) in 1% NDS for 4 h at 37°C. Samples were mounted with Prolong Gold Antifade DAPI (Thermo Fisher, P36935), and z-stack images were acquired using confocal microscopy (Zeiss LSM 800 AxioObserver.Z1/7) using a 488 nm laser (excitation) and 517 nm long pass filter (emission) and a Plan-Apochromat 63/1.4 NA oil immersion objective. Secondary antibody controls were used to verify the lack of non-specific staining. A total of 4 tubes/animal from five animals were analyzed. Z-stacks containing 14-16 slices each at 150nm were obtained, maximum projections were generated, and the mean fluorescence intensity (MFI) of each endothelial tube was measured using ImageJ and averaged to determine a mean fluorescence value for each animal.

For the HAoECs, ECs were exposed to shear stress (6 dynes/cm^2^ or 20 dyne/cm^2^) or static conditions for 24 hours using a parallel flow chamber prior to fixation with 4% PFA. HAoECs were blocked with 5% NDS and treated with ABCA1 primary antibody (1:100) (Thermo Fisher, PA1-16789) or CSE (1:100 Proteintech, 12217-1-AP) overnight at 4 °C. HAoECs were then blocked with 5% NDS for 1 hour at 37 °C and incubated with donkey-α-rabbit 488 (1:1000) in 1% NDS for 4 hours at 37 °C. Samples were mounted with Prolong Gold Antifade DAPI. Images contained 8-10 images with 7-10 cells per image for three independent runs, and the MFI was determined using ImageJ. The LXR agonist T0901317 (100 nM) is a well-established inducer of ABCA1 expression and was used to validate the specificity of the ABCA1 antibody (Fig. 6).

### Application of SS to Mesenteric Arteries

Mesenteric conduit arteries (>300 μm) were cannulated and pressurized to 65mmHg using a gravity-based column system. Luminal flow was generated by adjusting the column heights [P_arterial_ ∼130mmHg and P_venous_ ∼35mmHg (HSS)]. Arteries were continuously superfused with HPSS (5 ml/min, 37°C). Mean pressure was maintained at 65 mmHg. Vessels were exposed to flow for 6 hours, followed by a viability response (Ach-dilation), before fixation with 4% PFA for 20 minutes while being continually pressurized. Vessels were then cut along their longitudinal axis, secured *en face,* and blocked with 5% NDS for 1 hour at room temperature. Vessel segments were then incubated overnight at 4 °C with ABCA1 primary antibody (1:100 – PA1-16789, Thermo) in 1% NDS in phosphate-buffered saline. Secondary antibody AF-488 (1:1000, Jackson Lab) was applied for 1 hour at room temperature in 1% NDS in phosphate-buffered saline and mounted with Prolong Gold. Five images from various parts of the artery were captured per condition. ABCA1 intensity was measured using ImageJ.

### RT-qPCR Studies

HAoECs were exposed to either static, 6 dynes/cm^2^, or 20 dynes/cm^2^ conditions for 24 hours. In a separate set of experiments, HAoECs were treated with methyl-β-cyclodextrin (MβCD, 1mM) to deplete membrane cholesterol or MβCD + cholesterol (MβCD-Chol, 20mM) to increase membrane cholesterol content for 24 hours. RNA was isolated using an RNA extraction kit (Macherey-Nagel, 740955.50) according to the manufacture’s recommendations. cDNA was generated with 0.1 μg of RNA using the High-Capacity cDNA Reverse Transcription Kit (Thermo, 4374967). qPCR was performed using the TaqMan Fast Advanced Master Mix (Thermo 4444556). TaqMan Gene Expression Assay Primers for ABCA1 (4331182) and GAPDH (4331182) were purchased from Thermofisher. GAPDH was used as a housekeeping gene to normalize the expression of ABCA1. Data was normalized using the 2-delta CT method.

### Vasodilation Studies

The mesenteric arcade was removed and immediately placed in ice-cold HPSS. After the removal of veins and adipose tissue, third- to fourth-order mesenteric arteries (with an inner diameter of 100–200 μm) were isolated and cannulated with glass micropipettes, and gently flushed to remove blood from the lumen, as previously described^22,23^. The artery was stretched longitudinally to approximate its in situ length and pressurized with a servo-controlled peristaltic pump (Living Systems) to 75 mmHg. The vessel chamber was superfused with HPSS at 5 ml/min at 37°C and allowed to equilibrate for 15 min. To determine the contribution of ABCA1 on the H_2_S-mediated vasodilatory response, arteries were pretreated luminally with the specific ABCA1 antagonist, probucol (100 μM), for one hour before arteries were pre-constricted to ∼30–50% with the thromboxane A_2_ analog, U-46619 (Cayman Chemical). To assess H_2_S-mediated vasodilation, arteries were treated with sodium hydrosulfide (NaHS) at concentrations of 1, 10, and 30 μM (Cayman Chemical). Images were obtained using an Eclipse TS100 microscope (Nikon) and CCD100M camera to measure ID using edge-detection software (IonOptix). Passive diameter was determined at the end of each experiment by superfusing the vessel in Ca^2+^-free HPSS HPSS containing: 130 NaCl, 4 KCl, 1.2 MgSO_4_, 3.6 EGTA, NaHCO_3_, 10 HEPES, 1.18 KH_2_PO_4_, and 6 glucose; pH adjusted to 7.4 with NaOH), containing 10 μM sodium nitroprusside and 0.1% bovine serum albumin (in mM) for 30 min. For all experiments, changes in vessel diameter (plateau response) are expressed as the percent reversal of U46619-induced vasoconstriction.

### Cholesterol efflux

HAoECs were cultured in the presence of probucol (100 μM) or vehicle for 24 hours. The media was then used to assess cholesterol efflux using Amplex-Red, according to the manufacturer’s instructions (Thermo Fisher A12216). A subset of experiments were also exposed to static or HSS conditions for 24 hrs.

### Statistics and Calculations

All data are expressed as means ± SD. Statistical comparisons were made using Prism 9 (GraphPad Software). The specific statistical tests used are reported in the figure legends. A probability of <0.05 was accepted as statistically significant for all comparisons. Percent dilation was calculated as (NaHS diameter – U46619 diameter) / (Calcium Free diameter – U46619 diameter) x100%.

## Results

### Effect of SS on ABCA1 expression and membrane cholesterol in HAoECs

We exposed HAoECs to a SS of 6 dynes/cm^2^, 20 dynes/cm^2^, or static conditions for 24 hrs. Application of SS increased protein ABCA1 expression in HAoEC (Fig. 1a,b). SS reduced Abca1 mRNA levels in HAoEC (Fig. 1c). Depletion of membrane cholesterol reduced ABCA1 mRNA levels. Conversely, supplementing membrane cholesterol increased ABCA1 mRNA levels (Fig. 1d). To validate that changes in ABCA1 were associated with reductions in membrane cholesterol, we employed filipin staining to demonstrate that exposing cells to SS reduces the cholesterol content of the EC membrane (Fig. 2).

**Figure 1.**
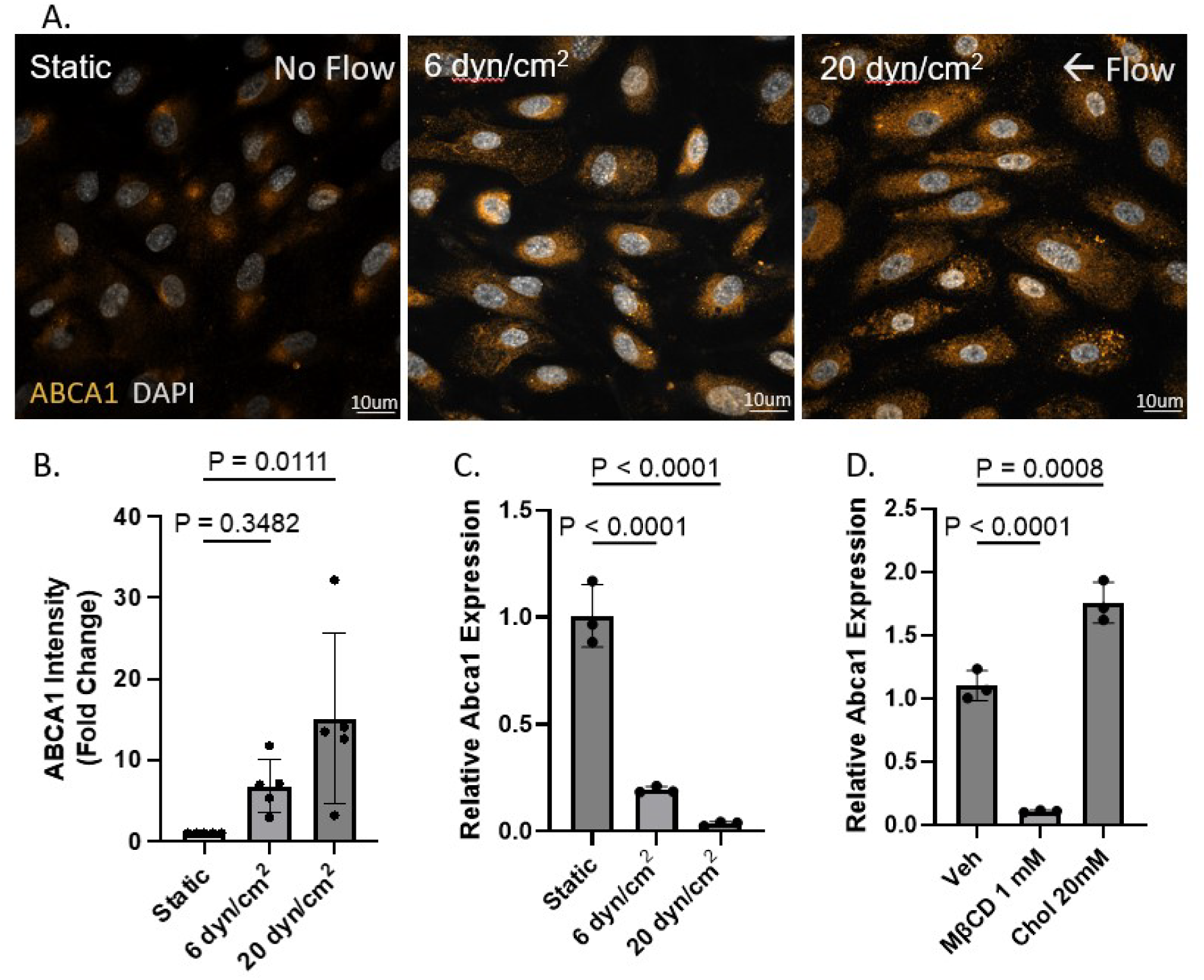
Shear stress increases ABCA1 expression. (A) Representative images of ABCA1 expression (yellow) in response to static, 6 dynes/cm^2^, or 20 dynes/cm^2^ in human aortic endothelial cells. (B) Quantification of mean fluorescence intensity for ABCA1 expression. 8-10 images were acquired per condition (n=5/group) with 6 – 7 cells per image. (C) ABCA1 mRNA expression normalized to GAPDH in response to static conditions, 6 dynes/cm^2^, or 20 dynes/cm^2^ in HAoECs. (D) ABCA mRNA in response to cholesterol manipulation in HAoECs. One-way ANOVA with Tukey’s Post-hoc test was used for statistical analysis.

**Figure 2.**
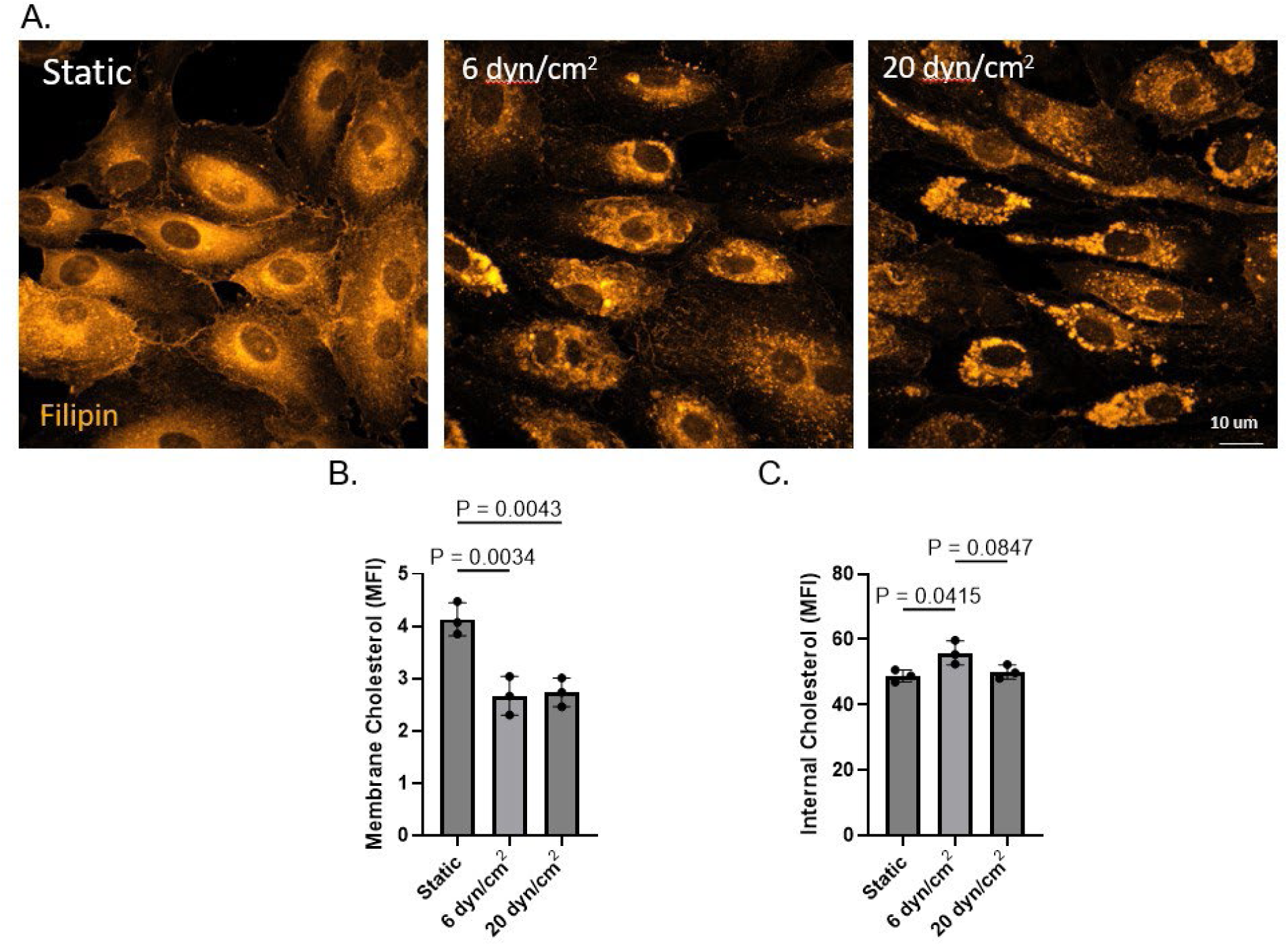
Shear stress decreases membrane cholesterol. (A) Representative images of membrane cholesterol expression in response to static, 6 dynes/cm^2^, or 20 dynes/cm^2^ in human aortic endothelial cells. (B) Quantification of mean fluorescence intensity for filipin III staining (orange) for the plasma membrane and (C) internal cholesterol content. 8-10 images per condition (n=3/group) with 6 – 7 cells per image were acquired. One-way ANOVA with Tukey’s Post-hoc test was used for statistical analysis.

### Effect of SS on ABCA1 expression in mesenteric conduit arteries

To assess the effect of SS on ABCA1 expression, pressurized mesenteric arteries were exposed to either 12 or 20 dynes/cm^2^ for 6 hours before fixation. SS elicited an increase in ABCA1 expression compared to arteries exposed to 12 dyn/cm^2^ SS (Fig.3). We have previously examined ABCA1 expression in freshly isolated paraformaldehyde-fixed endothelial cell tubes from both large and small mesenteric arteries.^24^ ABCA1 expression and membrane cholesterol were significantly lower in tubes from large arteries compared to small resistance arteries. ABCA1 expression appears greater in freshly isolated EC compared to arteries exposed to SS ex vivo. The difference in expression between the freshly isolated EC and arteries exposed to 6 hours of ex vivo flow may be due to differences in the absolute magnitude of shear stress in the two settings.

**Figure 3.**
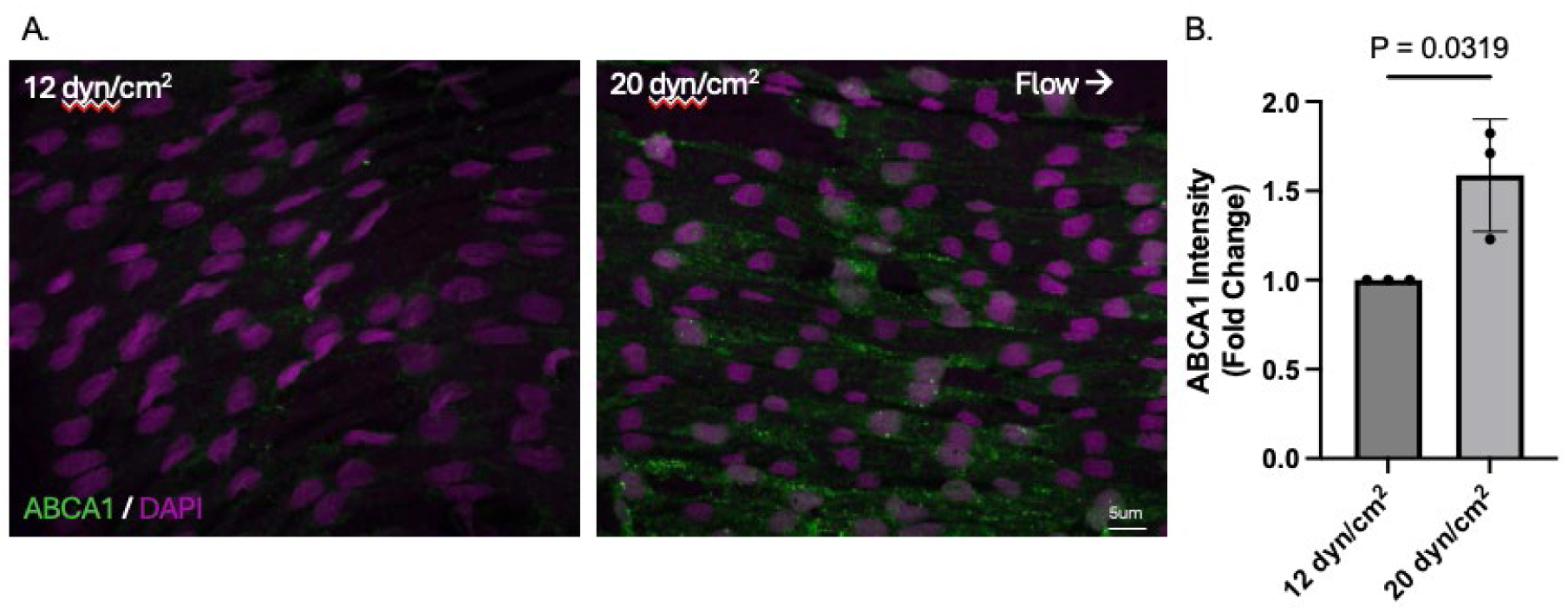
Increasing shear stress increases ABCA1 expression. (A) Representative images of ABCA1 expression in response to increasing shear stress in intact large mesenteric arteries. (B) Quantification of mean fluorescence intensity expressed as fold change for ABCA1 expression. Five images per artery were acquired per condition (n=3/group). Nuclear stain DAPI (magenta), ABCA1 (green). Student’s t-test was used for statistical analysis.

### Effect of ABCA1 inhibition on membrane cholesterol and H_2_S-mediated vasodilation in mesenteric resistance arteries

Inhibition of EC ABCA1 in small mesenteric arteries (<150 μm) increased EC membrane cholesterol and reduced cholesterol efflux from HAoECs (Fig. 4a-c). In addition, inhibition of ABCA1 significantly attenuated the SS-induced increase in cholesterol efflux (Fig 4d). Administration of increasing concentrations of H_2_S elicited a dose-dependent vasodilation in arteries treated with the vehicle. Inhibition of ABCA1 attenuated H_2_S-mediated vasodilation compared to control (Fig. 5).

**Figure 4.**
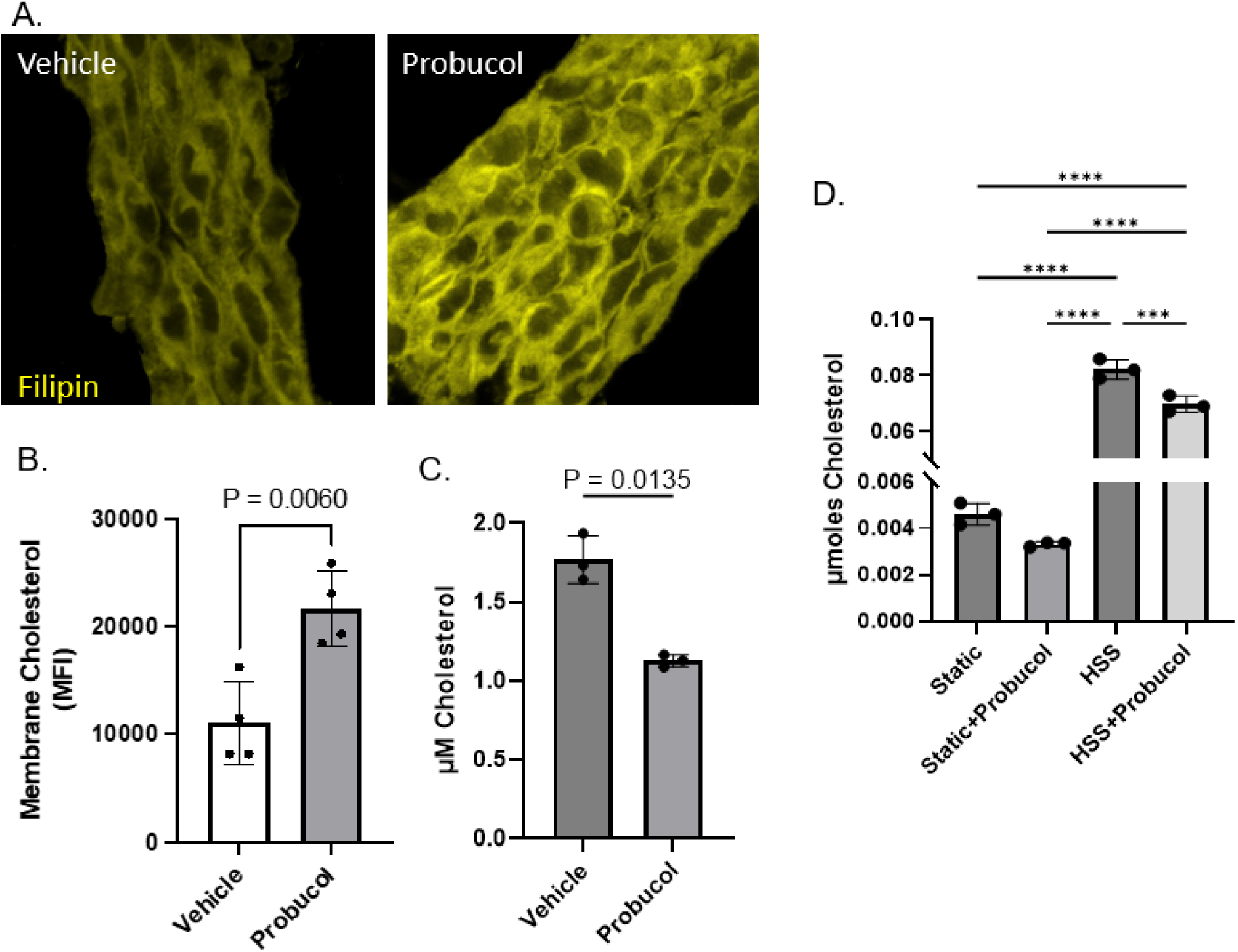
Inhibition of ABCA1 increases membrane cholesterol and reduces cholesterol efflux. (A) Representative images of membrane cholesterol in response to vehicle or probucol in endothelial tubes isolated from small mesenteric arteries (< 150 µm). (B) Quantification of mean fluorescence intensity for filipin III staining (yellow). 4 tubes/artery from 5 animals. (C) Quantification of cholesterol efflux from HAoECs. One-way Anova was used for statistical analysis. (D) Quantification of cholesterol efflux from HAoECs in response to shear stress. Student’s t-test or One-Way ANOVA with Tukey’s Post-hoc test was used for statistical analysis. **** P<0.0001, *** P< 0.0007.

**Figure 5.**
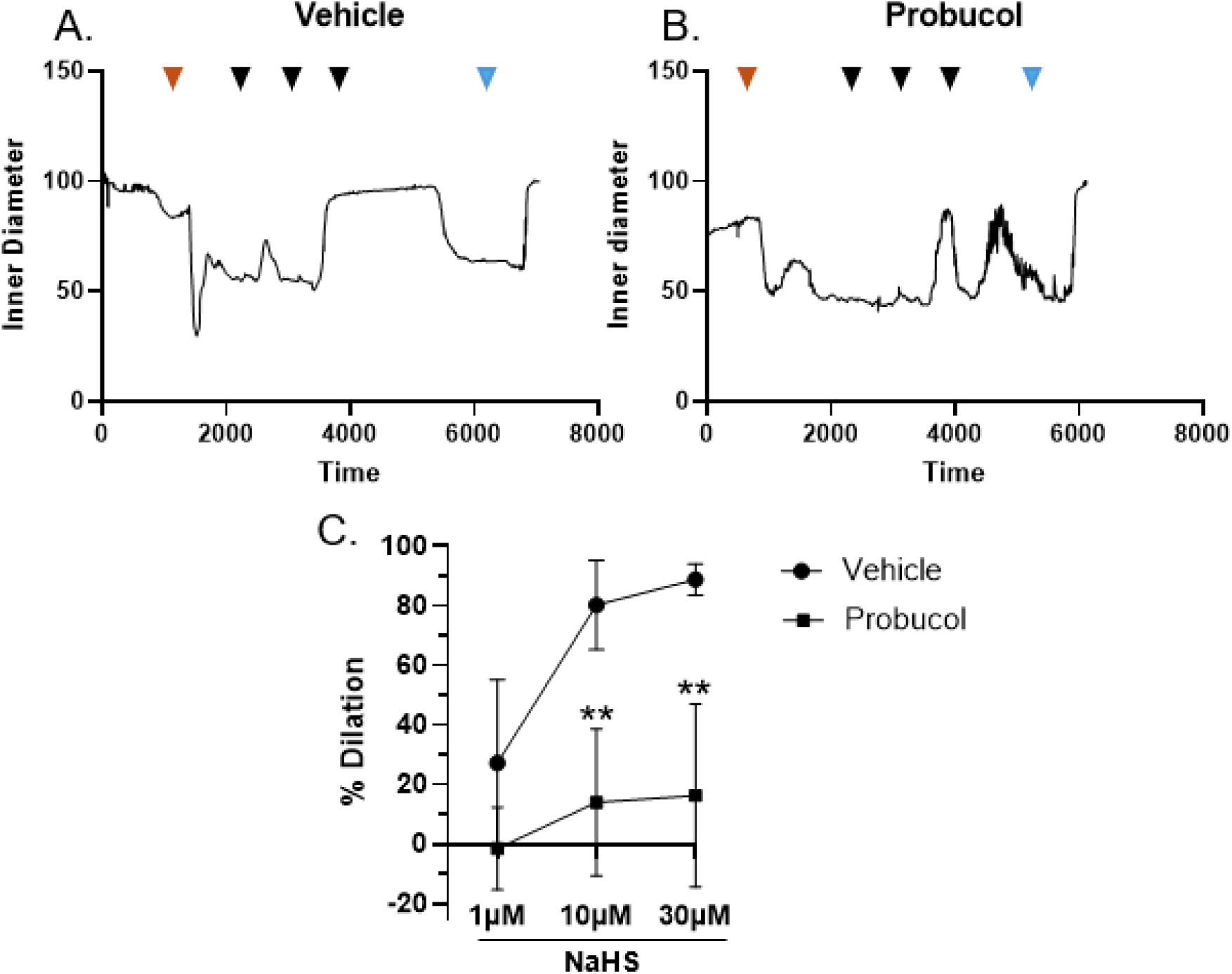
Inhibition of ABCA1 attenuates H_2_S-mediated vasodilation. (A) Representative trace of vehicle treated artery, The red arrow indicates pre-constrictor U46619, black arrows indicate H_2_S dosage, and the blue arrow indicates Ca^2+^ free PSS. (B) representative trace of probucol treated artery. (C) Summary responses of NaHS-dependent dilation in arteries treated with vehicle (n = 5) or the ABCa1 inhibitor probucol (100 µM, n = 5). Unpaired students t test was used for statistical analysis.

**Figure 6.**
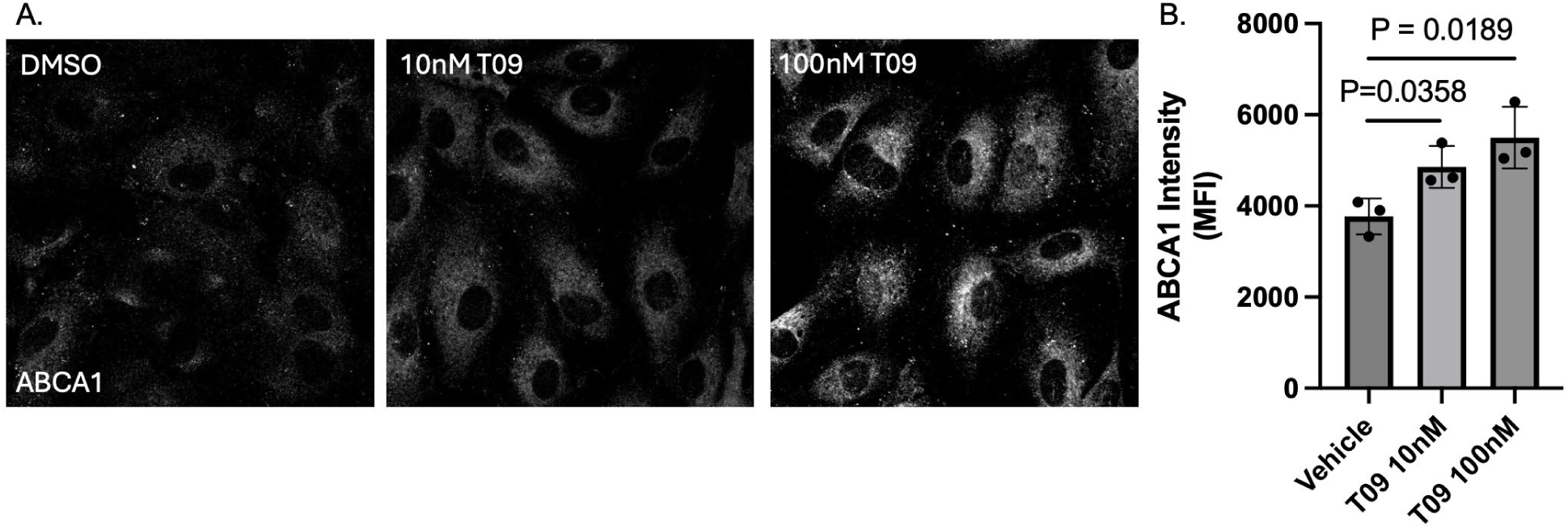
The liver X receptor agonist (T0901317) increases ABCA1 expression in HAoECs. (A) Representative images of ABCA1 expression in response to vehicle (DMSO) or T09 (10 µM and 100 µM). (B) Quantification of mean fluorescence intensity for ABCA1 expression. 8-10 images per condition (n=3) with 6 – 7 cells per image. One-way ANOVA with Tukey’s Post-hoc test was used for statistical analysis.

## Discussion

Endothelial function is crucial for maintaining vascular homeostasis, and disruptions in endothelial signaling pathways can lead to the development of cardiovascular disease. This study demonstrates a novel role for ABCA1 in regulating membrane cholesterol levels in endothelial cells. To our knowledge, this is the first study to demonstrate a role for ABCA1 in SS-mediated reductions in membrane cholesterol. The findings suggest that ABCA1 regulation of membrane cholesterol impacts hydrogen sulfide (H_2_S)-mediated vasodilation with potential implications for the regulation of microvascular function and disease pathophysiology. The key findings of this study are (1) SS increased ABCA1 expression in endothelial cells but reduced mRNA transcripts, (2) inhibition of ABCA1 increased membrane cholesterol levels and reduced cholesterol efflux, and (3) ABCA1 inhibition attenuated H_2_S-mediated vasodilation.

The regulation of membrane cholesterol by ABCA1 in resistance arteries has significant physiological implications. Resistance-sized arteries are critical regulators of vascular resistance and blood pressure, and their endothelial function is highly influenced by membrane cholesterol content^25,26^. Our previous work demonstrated that five up-regulated, differentially expressed genes involved in cholesterol transport (Nfkbia, Apoe, Lrp1, Abca1, Pltp) were present in ECs from small mesenteric arteries compared to ECs from large arteries. NF-kappa-B inhibitor alpha (*NFKBIA*) has been shown to enhance cholesterol transport in macrophages by inhibiting NFKB,^27,28^, whereas ApoE facilitates cholesterol transport through direct binding with ABCA1^29,30^. The low-density lipoprotein receptor-related protein (*LRP1*) has been shown to regulate lipid and glucose metabolism and is a known co-activator of a transcriptional regulator of ABCA1, PPAR gamma^31,32,33^. Moreover, phospholipid transfer protein (*PLTP*) functions as an auxiliary protein that facilitates ABCA1-mediated cholesterol transport^34^. This gene ontology pathway analysis related to cholesterol transport supports the postulate that ABCA1 is a key determinant of membrane cholesterol heterogeneity.

Our previous work has shown that H_2_S-mediated vasodilation in rat mesenteric arteries involves the activation of EC transient receptor potential cation subfamily V member 4 channel (TRPV4) channels and EC large conductance (BK_Ca_) Ca^2+^-activated potassium channels (Morin et al. 2022). TRPV4 and BK_ca_ contain cholesterol recognition/interaction amino acid consensus sequences (CRAC/CARC)^35,36^. Indeed, membrane cholesterol has been shown to negatively regulate TRPV4 and BK_Ca_ channel activity through direct protein-sterol interactions^35,37^. In addition to channel regulation via direct protein-sterol interactions, changes in membrane cholesterol may alter signaling microdomains. We have previously demonstrated that the depletion of membrane cholesterol is required to demonstrate a role for TPRV4 channels in muscarinic receptor-mediated vasodilation in rat gracilis arteries (Morin et al., 2022). In addition, we have demonstrated that depletion of membrane cholesterol with MbCD in arteries with higher cholesterol content (large mesenteric arteries) enhances H_2_S dilation. Conversely, loading arteries with MBCD-chol in arteries with lower native cholesterol content (small arteries) abolished H_2_S-mediated vasodilation. Interestingly, loading these arteries with the cholesterol enantiomer, epicholesterol, did not affect H_2_S vasodilation in these arteries. Suggesting that the effect of cholesterol on H_2_S vasodilation is due to protein-sterol interactions rather than changes in physical properties of the membrane (e.g., membrane fluidity).^38^ Our current findings suggest that increases in membrane cholesterol through ABCA1 inhibition attenuates H_2_S-mediated vasodilation in rat mesenteric resistance arteries, and cholesterol efflux was reduced in HAoECs. Our findings suggest that ABCA1 regulates endothelial membrane cholesterol by reducing cholesterol and potentiating the dilatory effect of H_2_S.

SS has been recognized as a key determinant of endothelial phenotype influencing processes such as nitric oxide production^39^, cellular polarization^40^, inflammation^41,42^, and permeability^43^. A previous report demonstrated a potential mechanism for enhanced ABCA1 expression in response to SS. Zhu *et al.* observed increases in key transcription factors involved in lipid homeostasis and ABCA1 regulation in human umbilical vein endothelial cells exposed to 2 dynes/cm^2^ or 12 dynes/cm^2^ SS. In addition, their findings suggest that SS increases mRNA levels of the CYP27A gene, which is responsible for oxysterol production. Additionally, mRNA levels of LXRα, an oxysterol-activated transcriptional regulator of *Abca1,* were increased in response to SS^13^. In contrast, Li *et al.* demonstrated that SS (40 dynes/cm^2^) reduced ABCA1 expression in human brain endothelial cells^14^. These discrepancies may be due to cell type differences or the magnitude of the applied shear stress. Our data suggest that increases in SS increase the expression of ABCA1, indicating that cholesterol transport mechanisms are actively modulated in response to shear stress. We demonstrated that HAoECs exposed to SS exhibit reduced membrane cholesterol levels compared to cells under static conditions.

In addition, this decrease in plasma membrane cholesterol was associated with a shift in the localization of free cholesterol to an endosomal-like state and ABCA1 has been shown to reside on the cell surface as well as endosomal compartments^44^. Notably, the application of shear stress reduced ABCA1 mRNA transcript levels despite a significant increase in ABCA1 protein. The sterol regulatory element-binding protein2 (SREBP2) is a critical regulator of genes involved in cellular cholesterol homeostasis. Classically, a decrease in cellular cholesterol leads to the activation and translocation of SREBP2 to the nucleus, thereby promoting the expression of genes involved in cholesterol uptake and de novo synthesis and suppressing the expression of cholesterol efflux genes^45^. In the present study, we observed that SS increased ABCA1 protein expression, which is associated with decreased membrane cholesterol and Abca1 mRNA levels. In addition, we demonstrate that depletion of membrane cholesterol with MbCD decreased Abca1 mRNA. In contrast, cholesterol supplementation increased Abca1 mRNA levels (Fig. 1). Taken together, these results suggest that SS may stabilize ABCA1, promoting cholesterol efflux, and that the loss of membrane cholesterol results in a downregulates Abca1 mRNA. One consideration in our cell culture model could be a change in the degradation rate, such that higher flow rates may protect ABCA1 from calpain-mediated degradation. Several studies have demonstrated the transcriptional and translation regulation of ABCA1 in the presence of cholesterol-depleting agents such as MBCD or Apo-A1. Studies suggest activation of ABCA1 to flux cholesterol from the cell resulted in stabilization of the ABCA1 complex and reduction in transcripts. Arakawa et al. demonstrate that apoA-1 binding to ABCA1 during cholesterol efflux protects the protein from degradation, increasing protein levels, which is associated with a decrease in ABCA1 transcript levels^46^. Over time, in response to continuous SS, a new set point for cholesterol homeostasis would be established, balancing cholesterol efflux, uptake, and synthesis^47^.

This study provides evidence that ABCA1 mediates cholesterol transport and regulates membrane cholesterol levels in endothelial cells, and that reductions in membrane cholesterol content facilitate ion channel activity, contributing to H_2_S signaling and promoting vasodilation. Taken together, these findings provide compelling evidence for a novel role for ABCA1 in vascular endothelial function. Our data also show that ABCA1 is a key mediator of endothelial membrane cholesterol in endothelial cells and that shear stress influences the expression of ABCA1, thereby regulating membrane cholesterol levels.

## Author contributions

J.R.A., L.V.G.B., N.L.K., and J.S.N. conceived and designed research; J.R.A. performed experiments; J.R.A. analyzed data; J.R.A., L.V.G.B., N.L.K., and J.S.N. interpreted results of experiments; J.R.A. and J.S.N. prepared figures; J.R.A. drafted manuscript; J.R.A., L.V.G.B., N.L.K., and J.S.N. edited and revised manuscript; J.R.A., L.V.G.B., N.L.K., and J.S.N. approved final version of manuscript.

## Work cited

1. Boulanger, C. M. B. Endothelium. https://www.ahajournals.org/doi/epub/10.1161/ATVBAHA.116.306940 (2016) doi:10.1161/ATVBAHA.116.306940.

2. Michiels, C. Endothelial cell functions. Journal of Cellular Physiology 196, 430–443 (2003).

3. Vascular Effects of Hydrogen Sulfide: Methods and Protocols. vol. 2007 (Springer New York, New York, NY, 2019).

4. Greaney, J. L. et al. Impaired Hydrogen Sulfide–Mediated Vasodilation Contributes to Microvascular Endothelial Dysfunction in Hypertensive Adults. Hypertension 69, 902–909 (2017).

5. Yang, G. et al. Hydrogen Sulfide Protects Against Cellular Senescence *via S* - Sulfhydration of Keap1 and Activation of Nrf2. Antioxidants & Redox Signaling 18, 1906–1919 (2013).

6. Kanagy, N. L., Szabo, C. & Papapetropoulos, A. Vascular biology of hydrogen sulfide. American Journal of Physiology-Cell Physiology 312, C537–C549 (2017).

7. Lv, B. et al. Hydrogen sulfide and vascular regulation – An update. J Adv Res 27, 85–97 (2020).

8. Mendiola, P. J., Morin, E. E., Gonzalez Bosc, L. V., Naik, J. S. & Kanagy, N. L. Role of Cholesterol in the Regulation of Hydrogen Sulfide Signaling within the Vascular Endothelium. Antioxidants (Basel) 11, 1680 (2022).

9. Das, A., Brown, M. S., Anderson, D. D., Goldstein, J. L. & Radhakrishnan, A. Three pools of plasma membrane cholesterol and their relation to cholesterol homeostasis. eLife 3, e02882 (2014).

10. Anderson, J. R. et al. Single-cell transcriptomic heterogeneity between conduit and resistance mesenteric arteries in rats. Physiological Genomics 55, 179–193 (2023).

11. Vasiliou, V., Vasiliou, K. & Nebert, D. W. Human ATP-binding cassette (ABC) transporter family. Hum Genomics 3, 281–290 (2009).

12. Oram, J. F. & Heinecke, J. W. ATP-Binding Cassette Transporter A1: A Cell Cholesterol Exporter That Protects Against Cardiovascular Disease. Physiological Reviews 85, 1343–1372 (2005).

13. Zhu, M. et al. Laminar Shear Stress Regulates Liver X Receptor in Vascular Endothelial Cells. *Arteriosclerosis*, Thrombosis, and Vascular Biology 28, 527–533 (2008).

14. Li, Z. et al. Cholesterol efflux regulator ABCA1 exerts protective role against high shear stress-induced injury of HBMECs via regulating PI3K/Akt/eNOS signaling. BMC Neuroscience 23, 61 (2022).

15. Vion, A.-C. et al. Shear Stress Regulates Endothelial Microparticle Release. Circulation Research 112, 1323–1333 (2013).

16. Roux, E., Bougaran, P., Dufourcq, P. & Couffinhal, T. Fluid Shear Stress Sensing by the Endothelial Layer. Frontiers in Physiology 11, (2020).

17. Lu, D. & Kassab, G. S. Role of shear stress and stretch in vascular mechanobiology. J. R. Soc. Interface. 8, 1379–1385 (2011).

18. Davis, M. J., Earley, S., Li, Y.-S. & Chien, S. Vascular mechanotransduction. Physiological Reviews 103, 1247–1421 (2023).

19. Naik, J. S., Osmond, J. M., Walker, B. R. & Kanagy, N. L. Hydrogen sulfide-induced vasodilation mediated by endothelial TRPV4 channels. Am J Physiol Heart Circ Physiol 311, H1437–H1444 (2016).

20. Rasband, W.S. (1997-2015) ImageJ. National Institutes of Health, Bethesda, Maryland, USA. - References - Scientific Research Publishing. https://www.scirp.org/reference/referencespapers?referenceid=1690059.

21. Wenceslau, C. F. et al. Guidelines for the measurement of vascular function and structure in isolated arteries and veins. American Journal of Physiology-Heart and Circulatory Physiology 321, H77–H111 (2021).

22. Garcia, S. M., Naik, J. S., Resta, T. C. & Jernigan, N. L. Acid-sensing ion channel 1a activates IKCa/SKCa channels and contributes to endothelium-dependent dilation. Journal of General Physiology 155, e202213173 (2022).

23. Morin, E. E., Salbato, S., Walker, B. R. & Naik, J. S. Endothelial cell membrane cholesterol content regulates the contribution of TRPV4 channels in ACh-induced vasodilation in rat gracilis arteries. Microcirculation 29, e12774 (2022).

24. Anderson, J. R. et al. Single-cell transcriptomic heterogeneity between conduit and resistance mesenteric arteries in rats. Physiol Genomics 55, 179–193 (2023).

25. Drożdż, D., Drożdż, M. & Wójcik, M. Endothelial dysfunction as a factor leading to arterial hypertension. Pediatr Nephrol 38, 2973–2985 (2023).

26. Beverley, K. M. & Levitan, I. Cholesterol regulation of mechanosensitive ion channels. Front. Cell Dev. Biol. 12, (2024).

27. Liu, K. et al. Inhibiting NF-B increases cholesterol efflux from THP-1 derived-foam cells treated with AngII via up-regulating the expression of ATP-binding cassette transporter A. Journal of Nanjing Medical University (2008).

28. Ferreira, V. et al. Macrophage-specific inhibition of NF-κB activation reduces foam-cell formation. Atherosclerosis 192, 283–290 (2007).

29. Krimbou, L. et al. Molecular interactions between apoE and ABCA1. Journal of Lipid Research 45, 839–848 (2004).

30. Getz, G. S. & Reardon, C. A. Apoprotein E and Reverse Cholesterol Transport. Int J Mol Sci 19, 3479 (2018).

31. Mao, H. et al. Endothelial LRP1 regulates metabolic responses by acting as a co-activator of PPARγ. Nat Commun 8, 14960 (2017).

32. Chawla, A. et al. A PPARγ-LXR-ABCA1 Pathway in Macrophages Is Involved in Cholesterol Efflux and Atherogenesis. Molecular Cell 7, 161–171 (2001).

33. Ghanbari-Niaki, A. et al. Heart ABCA1 and PPAR-α Genes Expression Responses in Male Rats: Effects of High Intensity Treadmill Running Training and Aqueous Extraction of Black Crataegus-Pentaegyna. Research in Cardiovascular Medicine 2, 153 (2013).

34. Oram, J. F., Wolfbauer, G., Vaughan, A. M., Tang, C. & Albers, J. J. Phospholipid Transfer Protein Interacts with and Stabilizes ATP-binding Cassette Transporter A1 and Enhances Cholesterol Efflux from Cells *. Journal of Biological Chemistry 278, 52379–52385 (2003).

35. Kumari, S. et al. Influence of membrane cholesterol in the molecular evolution and functional regulation of TRPV4. Biochemical and Biophysical Research Communications 456, 312–319 (2015).

36. Singh, A. K. et al. Multiple Cholesterol Recognition/Interaction Amino Acid Consensus (CRAC) Motifs in Cytosolic C Tail of Slo1 Subunit Determine Cholesterol Sensitivity of Ca2+- and Voltage-gated K+ (BK) Channels. J Biol Chem 287, 20509– 20521 (2012).

37. Wang, X.-L. et al. Caveolae Targeting and Regulation of Large Conductance Ca2+-activated K+ Channels in Vascular Endothelial Cells. Journal of Biological Chemistry 280, 11656–11664 (2005).

38. Mendiola, P. J., Morin, E. E., Gonzalez Bosc, L. V., Naik, J. S. & Kanagy, N. L. Role of Cholesterol in the Regulation of Hydrogen Sulfide Signaling within the Vascular Endothelium. Antioxidants (Basel) 11, 1680 (2022).

39. Sriram, K., Laughlin, J. G., Rangamani, P. & Tartakovsky, D. M. Shear-Induced Nitric Oxide Production by Endothelial Cells. Biophys J 111, 208–221 (2016).

40. Wojciak-Stothard, B. & Ridley, A. J. Shear stress–induced endothelial cell polarization is mediated by Rho and Rac but not Cdc42 or PI 3-kinases. J Cell Biol 161, 429–439 (2003).

41. Zakkar, M., Angelini, G. D. & Emanueli, C. Regulation of Vascular Endothelium Inflammatory Signalling by Shear Stress. http://www.eurekaselect.com.

42. Meng, Q. et al. Laminar shear stress inhibits inflammation by activating autophagy in human aortic endothelial cells through HMGB1 nuclear translocation. Commun Biol 5, 1–13 (2022).

43. Seebach, J. et al. Endothelial Barrier Function under Laminar Fluid Shear Stress. Lab Invest 80, 1819–1831 (2000).

44. Neufeld, E. B. et al. Cellular Localization and Trafficking of the Human ABCA1 Transporter * 210. Journal of Biological Chemistry 276, 27584–27590 (2001).

45. Zeng, L. et al. Sterol-responsive element-binding protein (SREBP) 2 down-regulates ATP-binding cassette transporter A1 in vascular endothelial cells: a novel role of SREBP in regulating cholesterol metabolism. J Biol Chem 279, 48801–48807 (2004).

46. Arakawa, R. & Yokoyama, S. Helical Apolipoproteins Stabilize ATP-binding Cassette Transporter A1 by Protecting It from Thiol Protease-mediated Degradation *. Journal of Biological Chemistry 277, 22426–22429 (2002).

47. Azuma, Y. et al. Retroendocytosis pathway of ABCA1/apoA-I contributes to HDL formation. Genes to Cells 14, 191–204 (2009).

